## Supplementary_Table3 for "Transcriptional Repression of *reaper* by Stand Still Safeguards Female Germline Development in *Drosophila*"

| Primer label | Primer sequence | Source |
| --- | --- | --- |
| stil_RT-PCR_Fw | TGAAGGTGGCTGCACTGGAAG | Current study |
| stil_ RT-PCR_Rv | CTGGCGCGGAGTCTAGTTGG | Current study |
| stil_qRT-PCR_Fw | TTGGGCAAACTCCATGAAATGA | Current study |
| stil_qRT-PCR_Rv | GTGGACATGCAAGCTGCTG | Current study |
| Tubulin_Fw | GGTAACCGTCGAAATCAGTGTT | (Xu et al., 2024) |
| Tubulin_Rv | TGGCTTTTCTGCTATACGTGTC | (Xu et al., 2024) |
| rp49_Fw | ATGACCATCCGCCCAGCATAC | (Xu et al., 2024) |
| rp49_Rv | CTGCATGAGCAGGACCTCCAG | (Xu et al., 2024) |
| sxl_Fw | CTCACCTTCGATCGAGGGTGTA | (Moschall et al., 2019) |
| sxl_Rv | GATGGCAGAGAATGGGAC | (Moschall et al., 2019) |
| rpr_Fw | CGATCAGGCGACTCTGTTG | Current study |
| rpr_Rv | CGGACTTTCTTCCGGTCTTC | Current study |
| hid_Fw | GAACTGCAGGAGCGAAAGC | Current study |
| hid_Rv | GTTCTTGTGTCCCGTCAACC | Current study |

**Supplementary Table 3. List of primers used for RT-PCR and qRT-PCR in this study.**

Moschall, R., Rass, M., Rossbach, O., Lehmann, G., Kullmann, L., Eichner, N., Strauss, D., Meister, G., Schneuwly, S., Krahn, M. P. & Medenbach, J. (2019). Drosophila Sister-of-Sex-lethal reinforces a male-specific gene expression pattern by controlling Sex-lethal alternative splicing. *Nucleic Acids Research*, *47*(5), 2276–2288. <https://doi.org/10.1093/nar/gky1284>

Xu, F., Suyama, R., Inada, T., Kawaguchi, S. & Kai, T. (2024). HemK2 functions for sufficient protein synthesis and RNA stability through eRF1 methylation during Drosophila oogenesis. *Development*, *151*(14). <https://doi.org/10.1242/dev.202795>
