## Supplementary figures and images for "Transcriptional Repression of *reaper* by Stand Still Safeguards Female Germline Development in *Drosophila*"

### Supplementary Figures

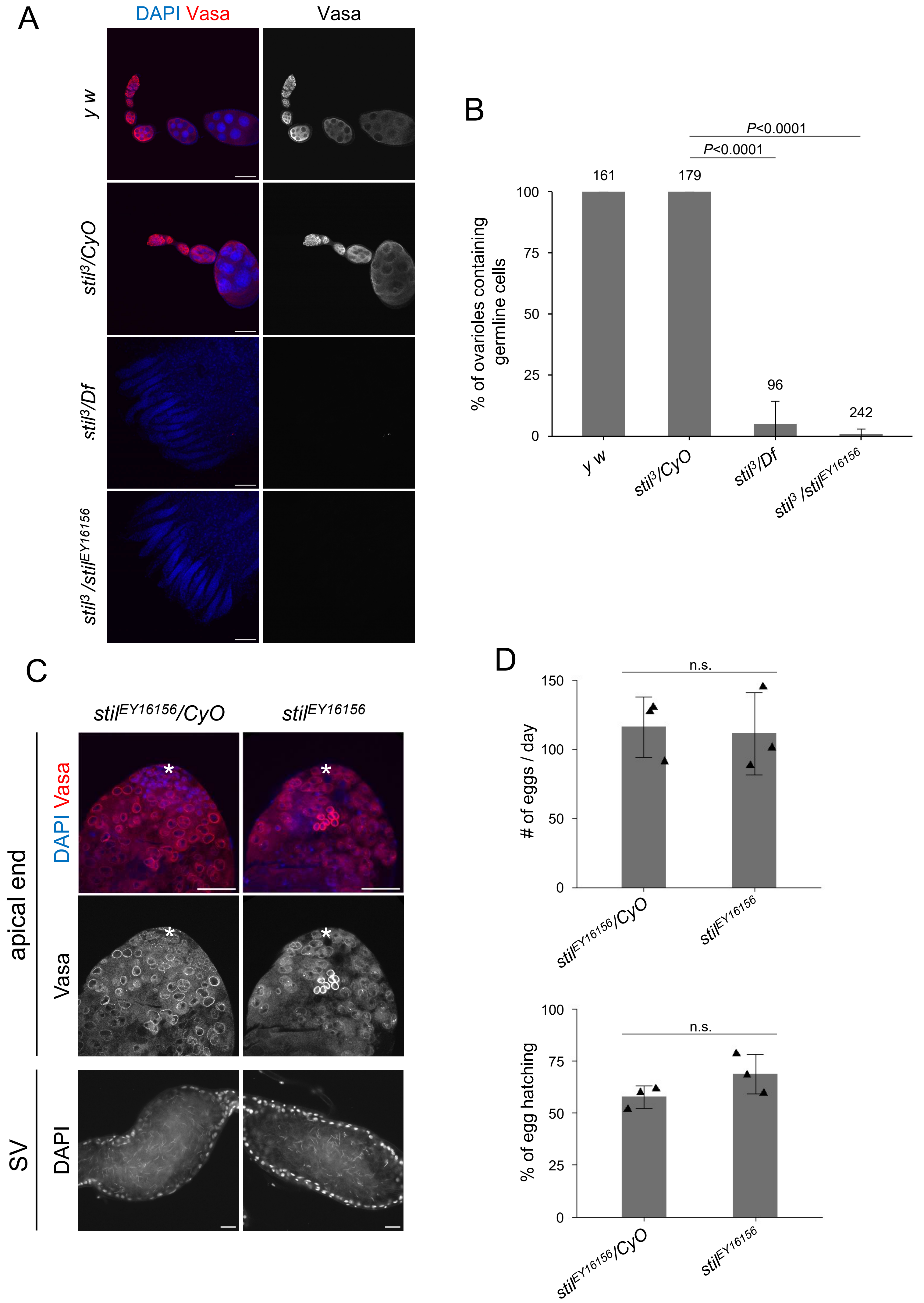

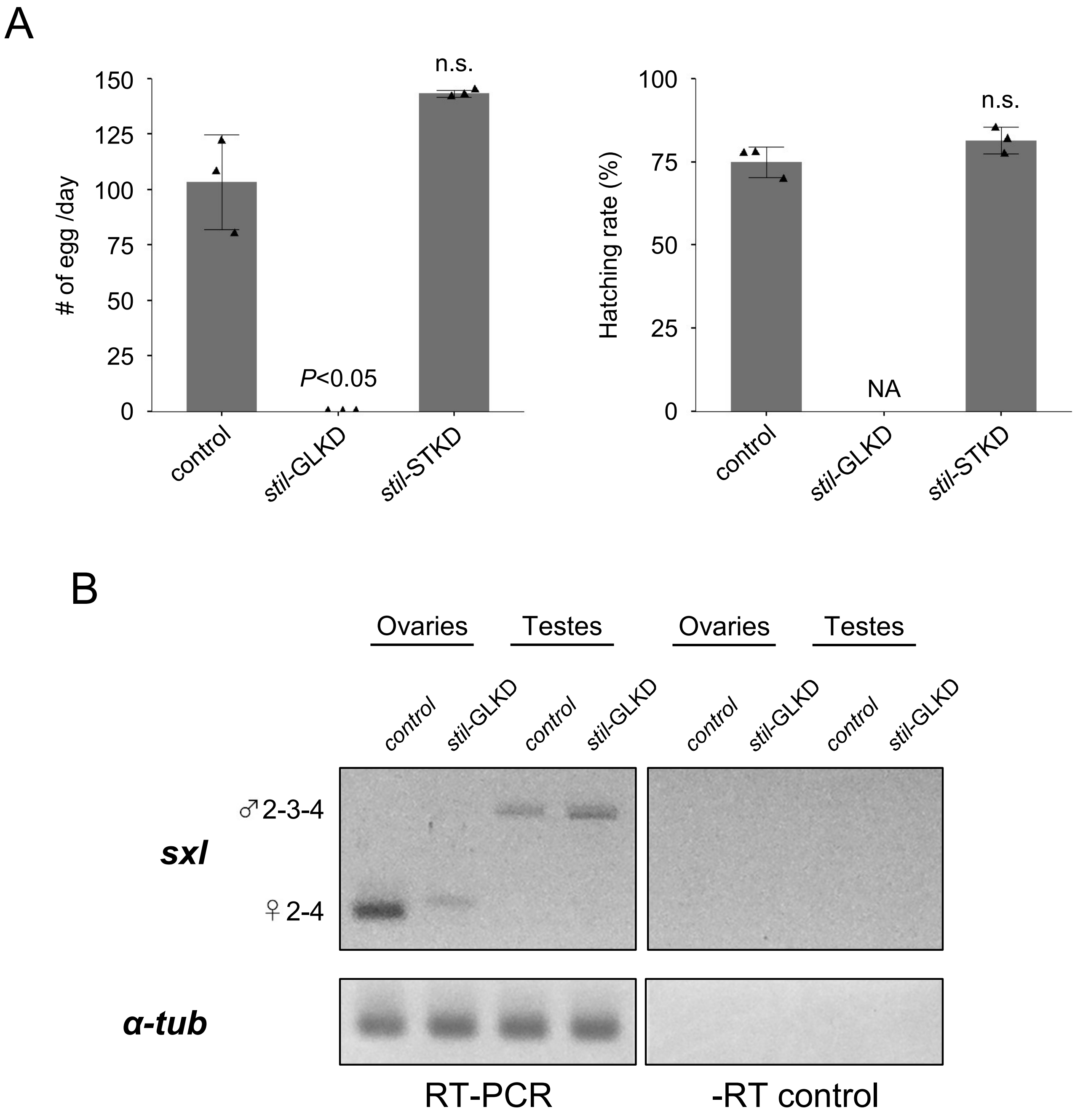

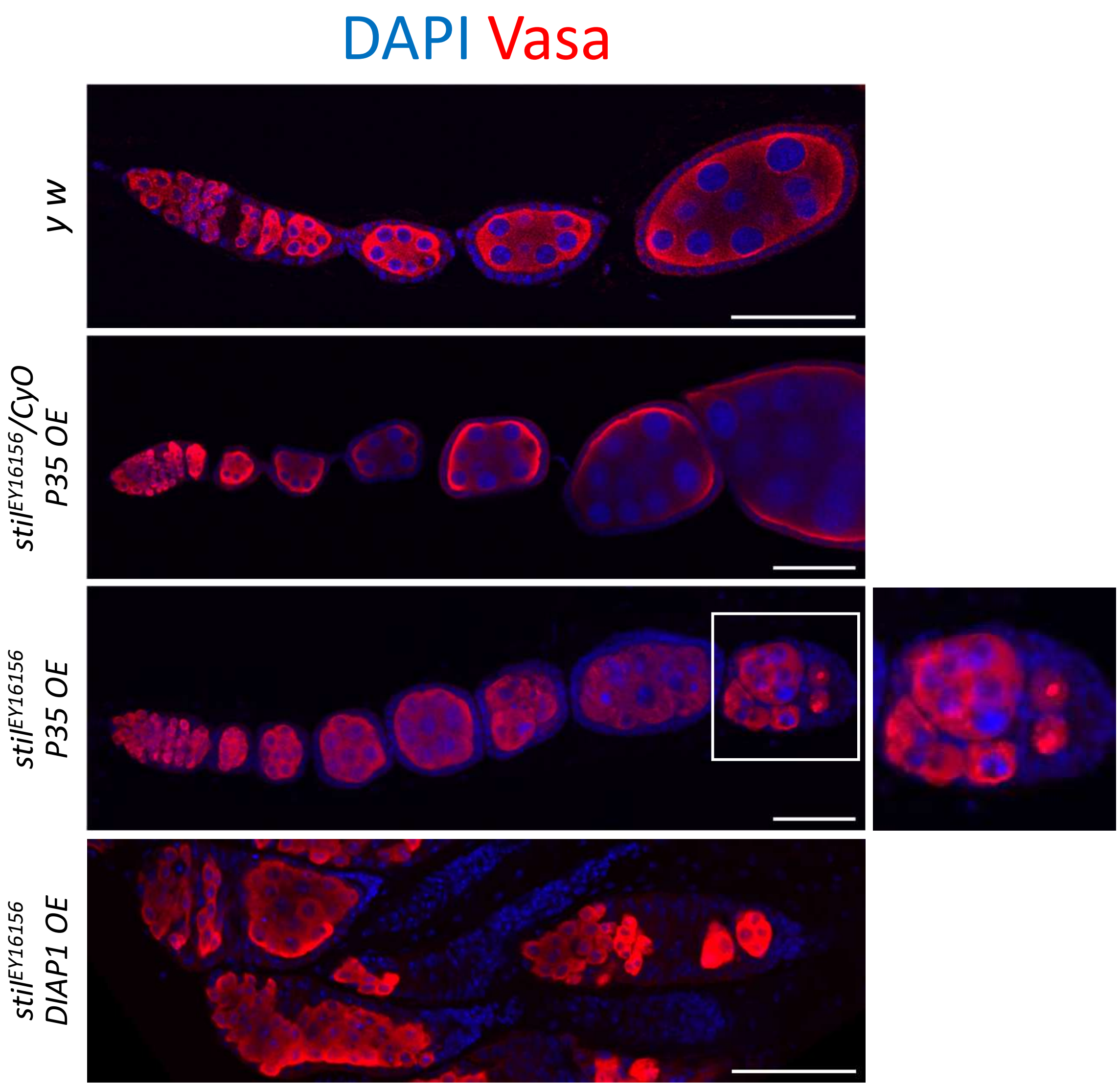

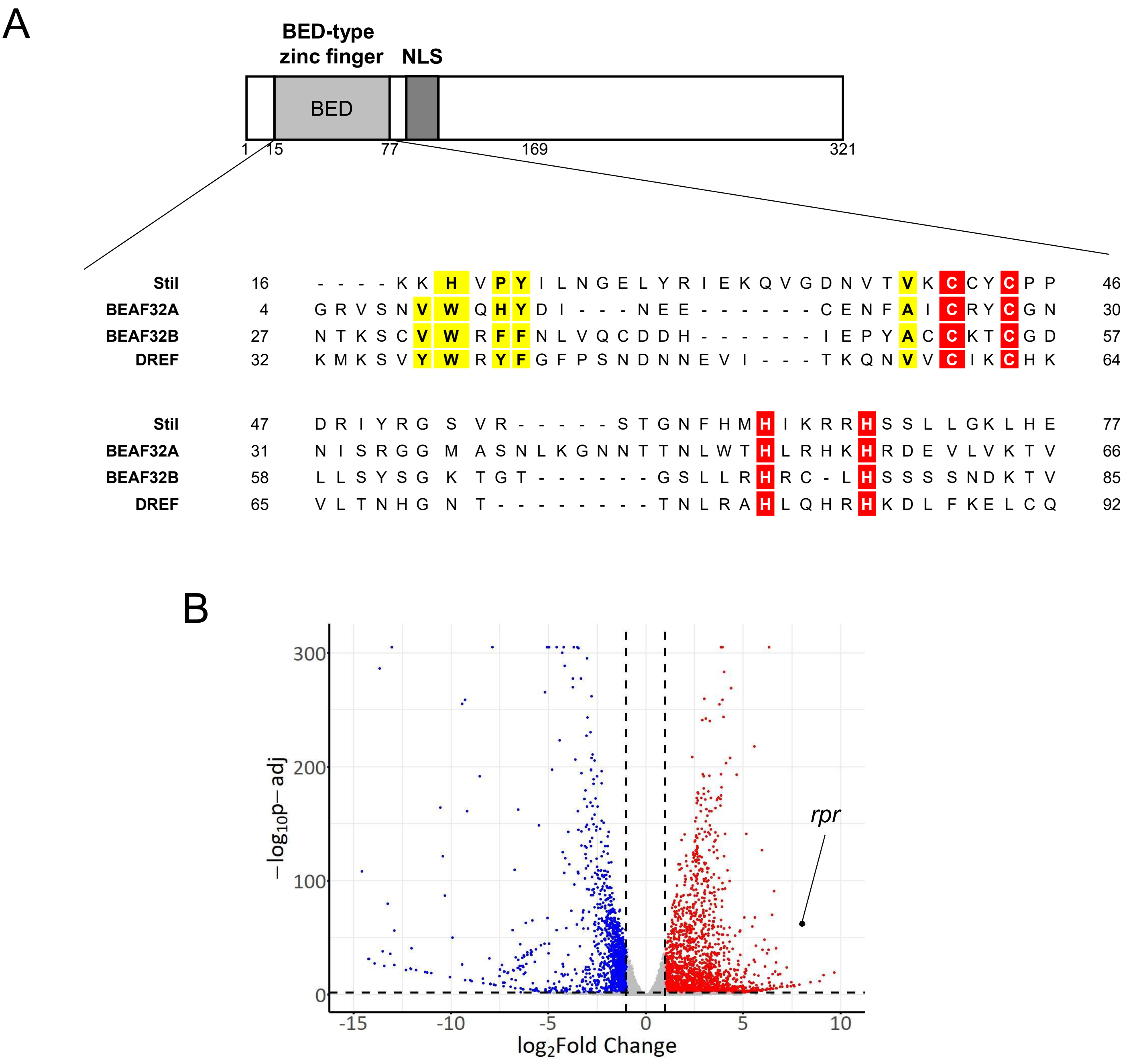

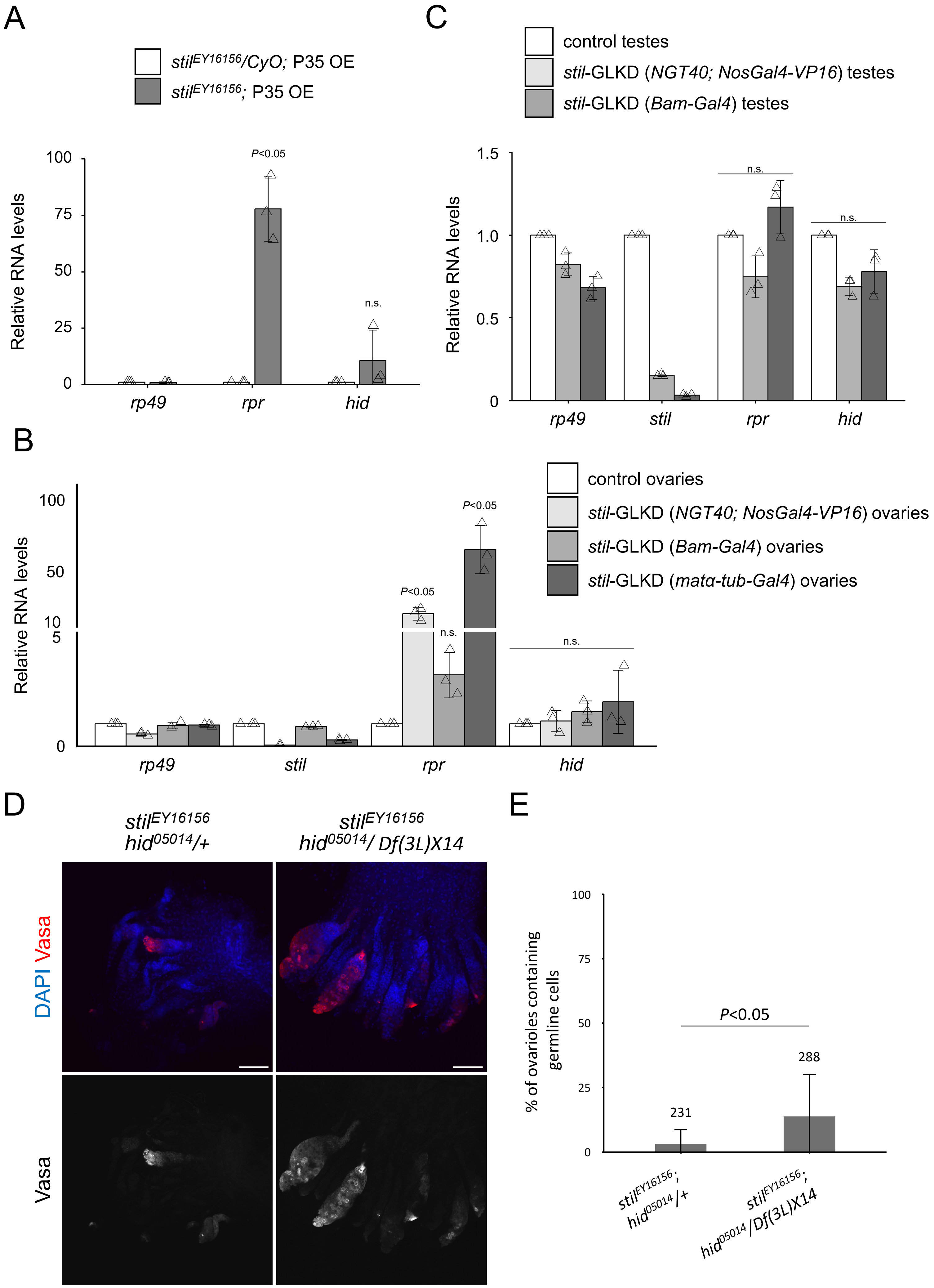

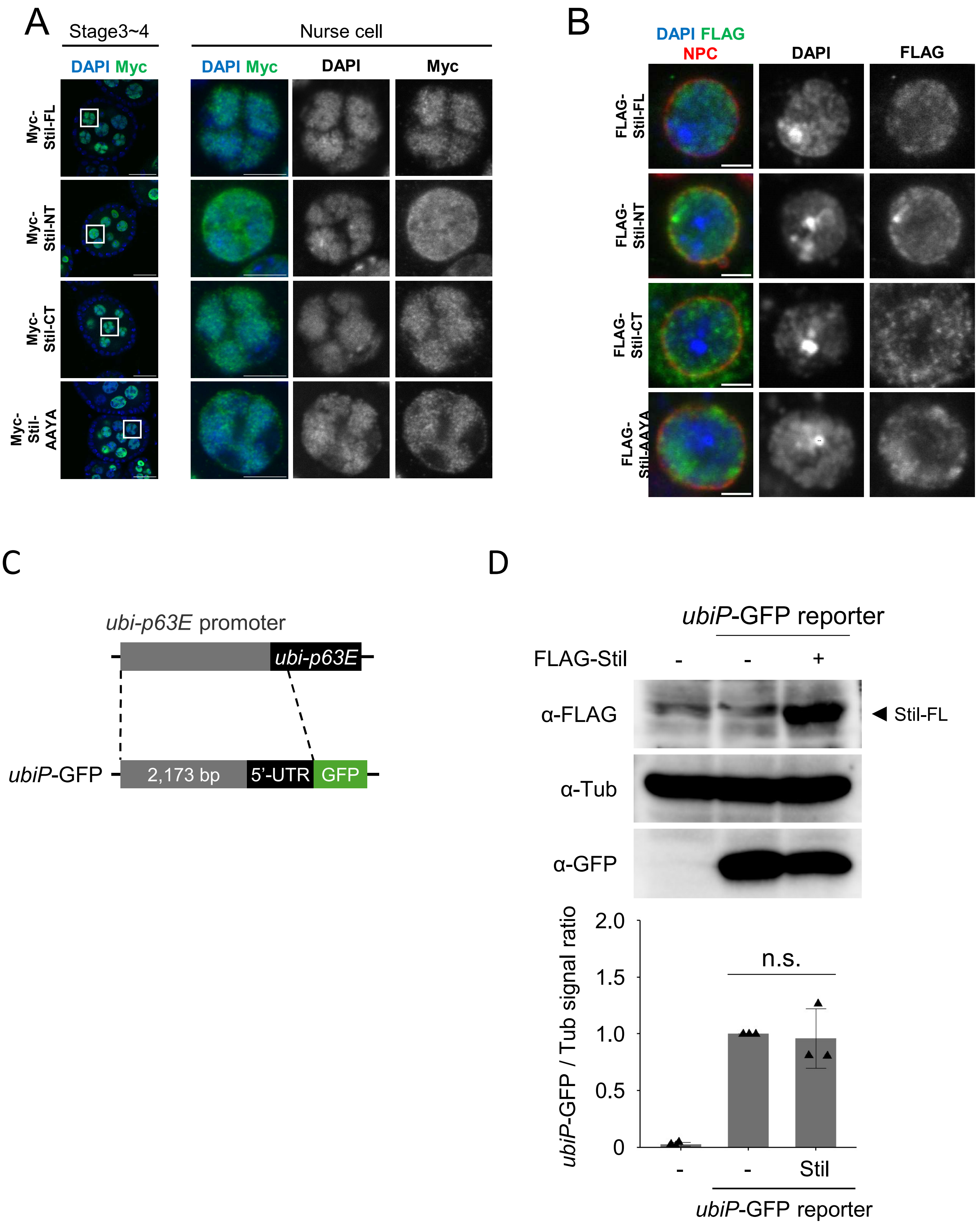
